# The utility of CRISPR activation as a platform to identify interferon stimulated genes with anti-viral function

**DOI:** 10.1101/2023.08.28.555046

**Authors:** Emily N. Kirby, Xavier B. Montin, Timothy P. Allen, Jaslan Densumite, Brooke N. Trowbridge, Michael R. Beard

**Affiliations:** Research Centre for Infectious Diseases; Department of Molecular and Biomedical Sciences, The University of Adelaide, Adelaide, South Australia, 5005, Australia; Department of Immunology, Faculty of Medicine, Siriraj Hospital, Mahidol University, Bangkok, Thailand

## Abstract

Interferon Stimulated Genes (ISGs) play key roles in the control of viral replication and dissemination. Understanding this dynamic relationship between the pathogen and host is critical to our understanding of viral life-cycles and development of potential novel anti-viral strategies. Traditionally, plasmid based exogenous prompter driven expression of ISGs has been used to investigate anti-viral ISG function, however there are deficiencies in this approach. To overcome this, we investigated the utility of CRISPR activation (CRISPRa), which allows for targeted transcriptional activation of a gene from its endogenous promoter. Using the CRISPRa-SAM system to induce targeted expression of a panel of anti-viral ISGs we showed robust induction of mRNA and protein expression. We then employed our CRISPRa-SAM ISG panel in several antiviral screen formats to test for the ability of ISGs to prevent viral induced cytopathic cell death (CPE) and replication of Dengue Virus (DENV), Zika Virus (ZIKV), West Nile Virus Kunjin (WNV_KUNV_), Hepatitis A Virus (HAV) and Human Coronavirus 229E (HCoV-229E). Our CRISPRa approach confirmed the anti-viral activity of ISGs like IFI6, IFNβ and IFNλ2 that prevented viral induced CPE, which was supported by high-content immunofluorescence imaging analysis. This work highlights CRISPRa as a rapid, agile, and powerful methodology to identify and characterise ISGs and viral restriction factors.

## Introduction

Viruses continue to pose a significant threat to human health, as highlighted by the continuing COVID-19 pandemic. Interferons and associated interferon stimulated genes (ISG) act a critical first line of defence in controlling viral replication. Activation of ISG responses is initiated by recognition of Pathogen Associated Molecular Patterns (PAMPs) by Pattern Recognition Receptors (PRRs) such as the Toll Like Receptors (TLRs) and cytosolic receptors RIG-I and MDA5 [1,2]. PRR activation initiates type I interferon (IFN, IFNα/IFNβ) expression that signals via the interferon receptor (IFNAR) to drive expression of over 300 ISGs (REF). These ISGs work to induce an anti-viral state in the cell either directly or indirectly, with the ultimate aim to control viral replication [3] (others). Therefore, it is important that the anti-viral potential of ISGs is well characterised as it will provide a greater understanding of the dynamic viral-host relationship, and insight for the development of novel anti-viral therapeutics.

Typically, characterisation of ISG activity utilises several techniques such as exogenous IFN stimulation or knockdown/knockout strategies such as interfering RNA (RNAi) [4–6] and CRISPR_KO_ [7–10] or ectopic plasmid based overexpression [11–14]. These methods have provided valuable insight into ISGs antiviral function, however they can be costly and labour intensive, creating an experimental bottleneck. Additionally, ectopic plasmid overexpression methods do not consider the limitations of excessive expression over and above physiological protein expression or splice variation, which is common for many ISGs [15]. To overcome these challenges, CRISPR activation (CRISPRa) has the potential to provide a suitable alternative. In contrast to traditional CRISPR_KO_ technology where Cas9 is catalytically active, leading to double stranded DNA cleavage (dsb), CRISPRa utilises a catalytically inactivate “dead” Cas9 (dCas9) that maintains DNA binding capacity [16–18]. When fused with transcriptional activators, dCas9 can be positioned by a single guide RNA (sgRNA) to the proximal promoter of a target gene where recruitment of additional transcriptional activators (e.g. histone demethylases), scaffolding proteins and ultimately RNA Pol, induces target gene transcription [18–20]. CRISPRa provides the benefit of inducing transcription directly from the endogenous promoter, limiting transcription to the physiological upper limit and expression of splice variants. Recently, CRISPRa genome-wide screening has become widely accessible, allowing the identification of both novel and canonical anti-viral host factors, however, very few have been ISGs [21–24]. It is possible that the anti-viral capacity of many ISGs may have been masked by stronger phenotypic responses, therefore, the utility of CRISPRa as tool to characterise ISG responses is to be determined.

In this study we determined the efficacy of 2 readily available systems, CRISPRa-SAM [17] and CRISPR-SunTag [16] at inducing transcriptional activation of a subset of ISGs in transient models without the need for IFN stimulation. As CRISPRa-SAM consistently achieved greater activation, CRISPRa ISG constitutively expressing cell lines were generated and utilised to determine an ISGs ability to inhibit viral replication by means of high-content immunofluorescence or luciferase assays. To further screen anti-viral activity we developed the cytopathic protection assay (CPA) to measure the rate of virus induced cytopathic cell death (CPE). Our arrayed screens confirmed the anti-viral activity of several well-established ISGs, such as IFI6 and IFNλ2, to inhibit viral induced CPE and replication of several viruses.

## Results

### CRISPRa, -SAM and -SunTag, induce transcriptional activation of targeted ISGs

We selected two widely available systems, CRISPRa-SAM (SAM) and CRISPRa-SunTag (SunTag), to validate transcriptional activation efficiency and determine which system would be appropriate for our anti-viral array screens. For both systems, dCas9 acts as a scaffold for interaction and/or fusion with several transcriptional activator proteins and an sgRNA (Fig 1). SAM incorporates a bacteriophage MS2 RNA aptamer sequence into the sgRNA that allows binding of the activator complex comprised of the bacteriophage MS2 coat protein and the transcription factors heat shock factor 1 (HSF1) and p65 (Fig 1A) [18] . To further enhance transcriptional activation, dCas9 is fused to VP64, a tetramer of the transcription factor VP16 derived from HSV-1. In contrast, SunTag utilises the “SunTag peptide”, 10 repeating units of the yeast master transcriptional regulator General Control Nondepressible 4 (GCN4), that is recognised by small chain variable fragment (ScFv) antibodies raised against GCN4 (Fig 1B) [25]. The ScFv_(GCN4)_ are fused with VP64 to initiate transcription and GFP to confirm ScFv expression. Together, these complexes recruit additional transcriptional activators (e.g. Oct-1, HCF, BRG1) that promote unwinding of DNA from chromatin, formation of the transcription scaffolding and subsequent RNA Pol binding [26–28].

**Figure 1:**
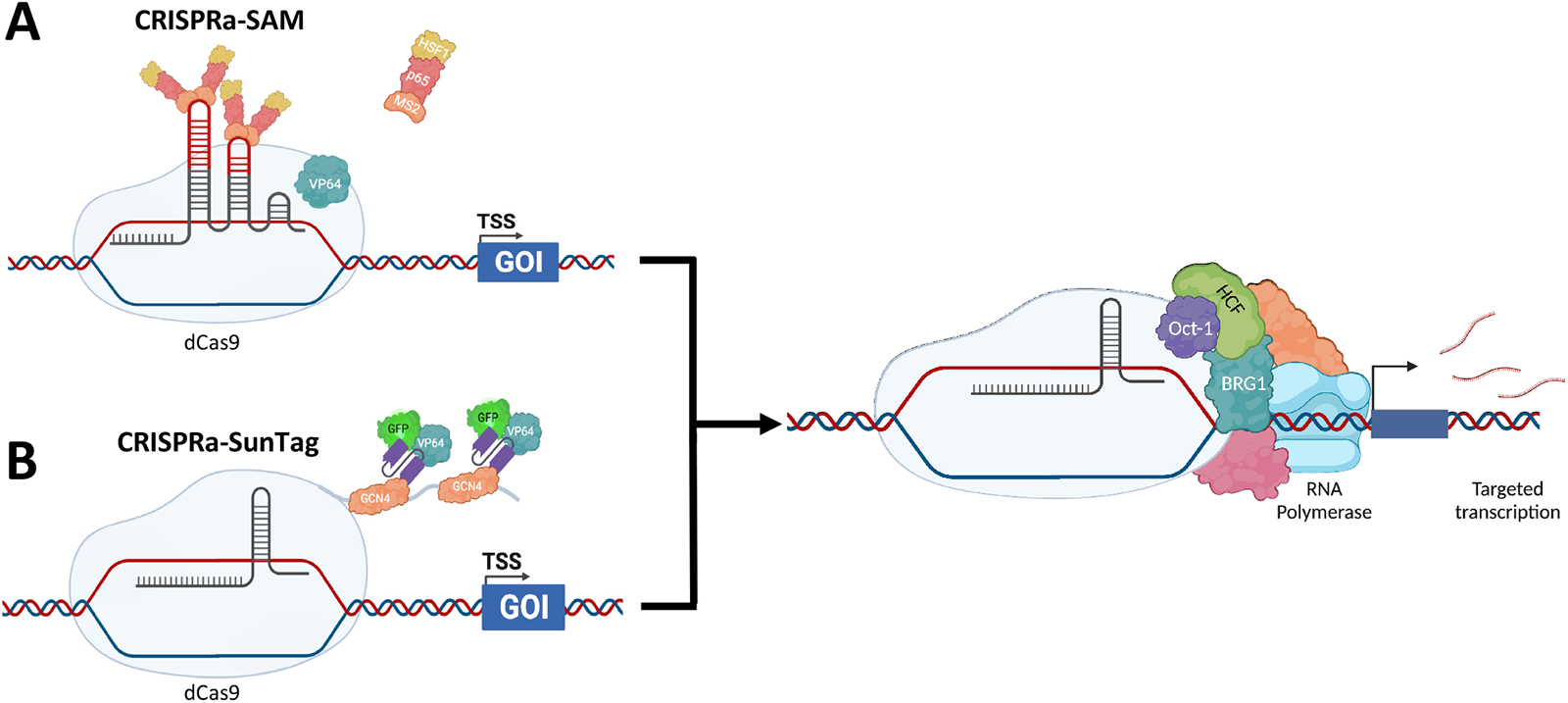
CRISPRa-SAM and CRISPRa-SunTag methods of inducing transcriptional activation. Currently, two of the most widely used CRISPRa systems are CRISPRa-SAM **(A)** and CRISPRa-SunTag **(B)**, which utilise differing transcriptional activators to activate transcription of a target gene by recruitment of additional transcription factors, transcription scaffolding proteins and ultimately RNA Pol.

Initially, we performed time-course experiments in HeLa cells to determine the level of transcriptional activation of the ISGs, IFI6 and IFITM1 by both SAM and SunTag systems. Cells were co-transfected with the respective SAM and SunTag vectors pXPR_502 or pCRISPRaV2, containing an sgRNA complementary for IFI6, IFITM1 or a non-target control (NTC) and the corresponding dCas9 and transcriptional activator vectors. qRT-PCR analysis indicated a time dependent increase of IFI6 and IFITM1 mRNA over a 72hr period for both SAM (Fig 2A) and SunTag (Fig 2B), however the SAM system produced consistently higher fold mRNA changes relative to time matched NTCs. This observation was supported by western blot of whole cell lysates from HeLas CRISPRa transfected with an IFITM1 specific or NTC sgRNA. We included a ΔdCas9 group where HeLa cells were transfected with the sgRNA, but not dCas9. This was to ensure that introduction of the CRISPRa systems does not inadvertently induce off-target ISG activation. IFITM1 was detected in the lysates of SAM transfected HeLas from 48hrs post-transfection, while IFITM1 was detected in SunTag transfected HeLas from 72hrs for SunTag (Fig 2C).

**Figure 2:**
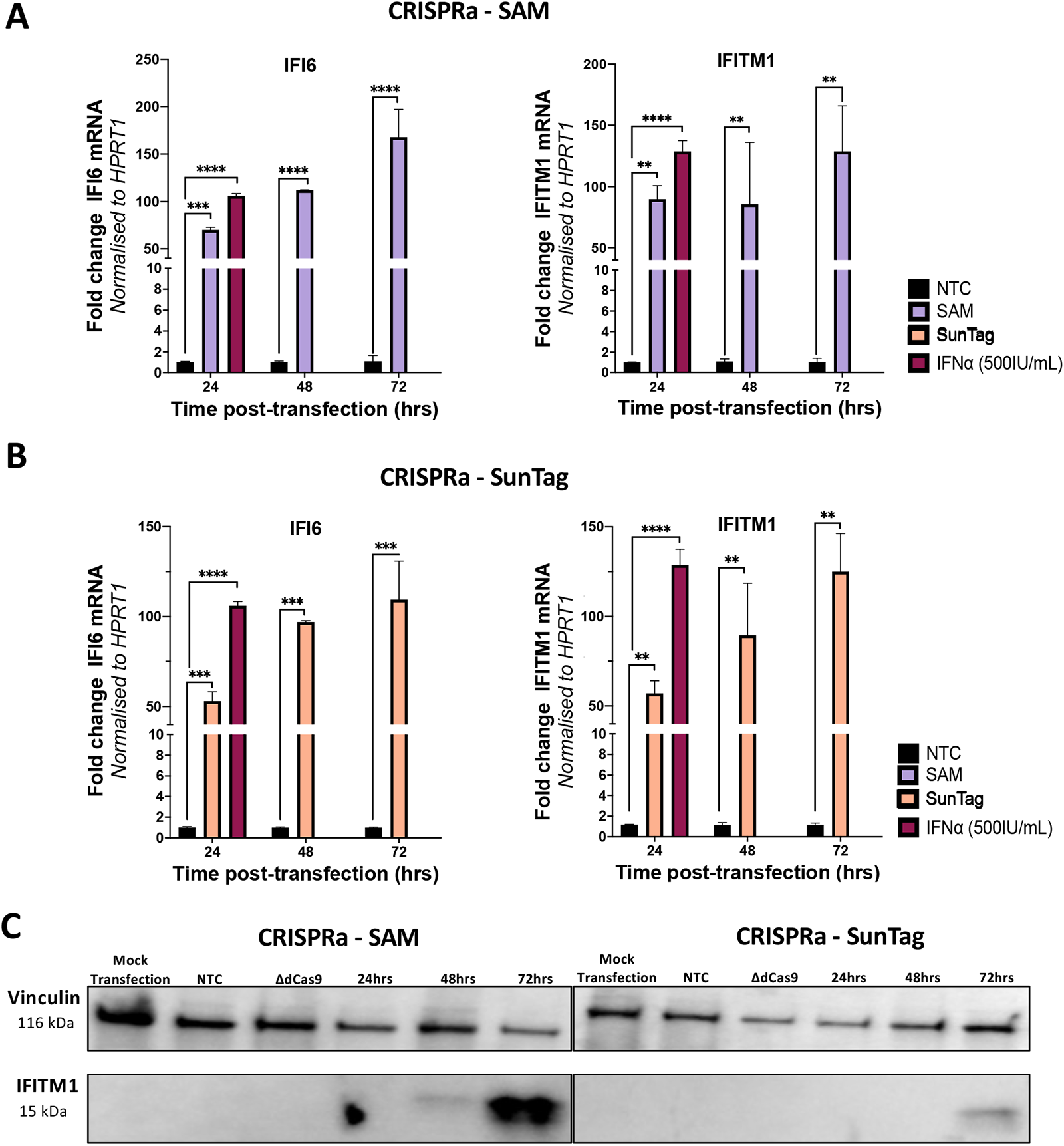
CRISPRa-SAM and SunTag induce transcriptional activation of target ISGs overtime. To determine the efficacy of the CRISPRa-SAM and SunTag, HeLas were transfected with the corresponding CRISPRa-SAM **(A)** or CRISPRa-SunTag **(B)** dCas9 and sgRNA plasmids for IFI6 or IFITM1 with total RNA extracted at the stated time points and qRT-PCR performed to determine CRISPRa mediated transcription of target ISGs relative to the non-targeting control (NTC) and IFN⍺ (500IU/mL) pre-treatment. Additionally for IFITM1, lysates were collected at stated timepoints and western blotting performed to confirm CRISPRa mediated transcriptional activation increase protein expression **(C)**. Statistical analysis was performed using Two-way ANOVA where * P ≤ 0.05, ** P ≤ 0.01, *** P ≤ 0.001, **** P ≤ 0.0001.

Given the diversity of *in vitro* culture lines, next we investigated if the efficacy of CRISPRa-SAM and -SunTag systems varied between different cell lines. HeLa and Huh7.5 cells were transfected with the respective dCas9 and either one of multiple distinct sgRNAs (termed sgRNA#1 and sgRNA #2) for the ISGs IFI6 and IFITM1. By qRT-PCR, we did not observe cell based significant differences in the ability of SAM (Fig 3A) or SunTag (Fig 3B) to induce transcription of IFI6 or IFITM1 mRNA, although SAM consistently induced greater activation. However, we did observe significant variation between the gene specific sgRNA themselves, where one sgRNA consistently outperformed the other. This is likely due to sgRNA design where well-established limitations, such as DNA accessibility and distance from the transcription start site alter activation efficacy [29]. Overall, SAM consistently outperformed SunTag, hence SAM was utilised for further screening experiments. In all CRISPRa ISG experiments we included stimulation of cells with IFNα, noting that in most cases the level of ISG induction was comparable between CRISPRa and IFNα stimulation.

**Figure 3:**
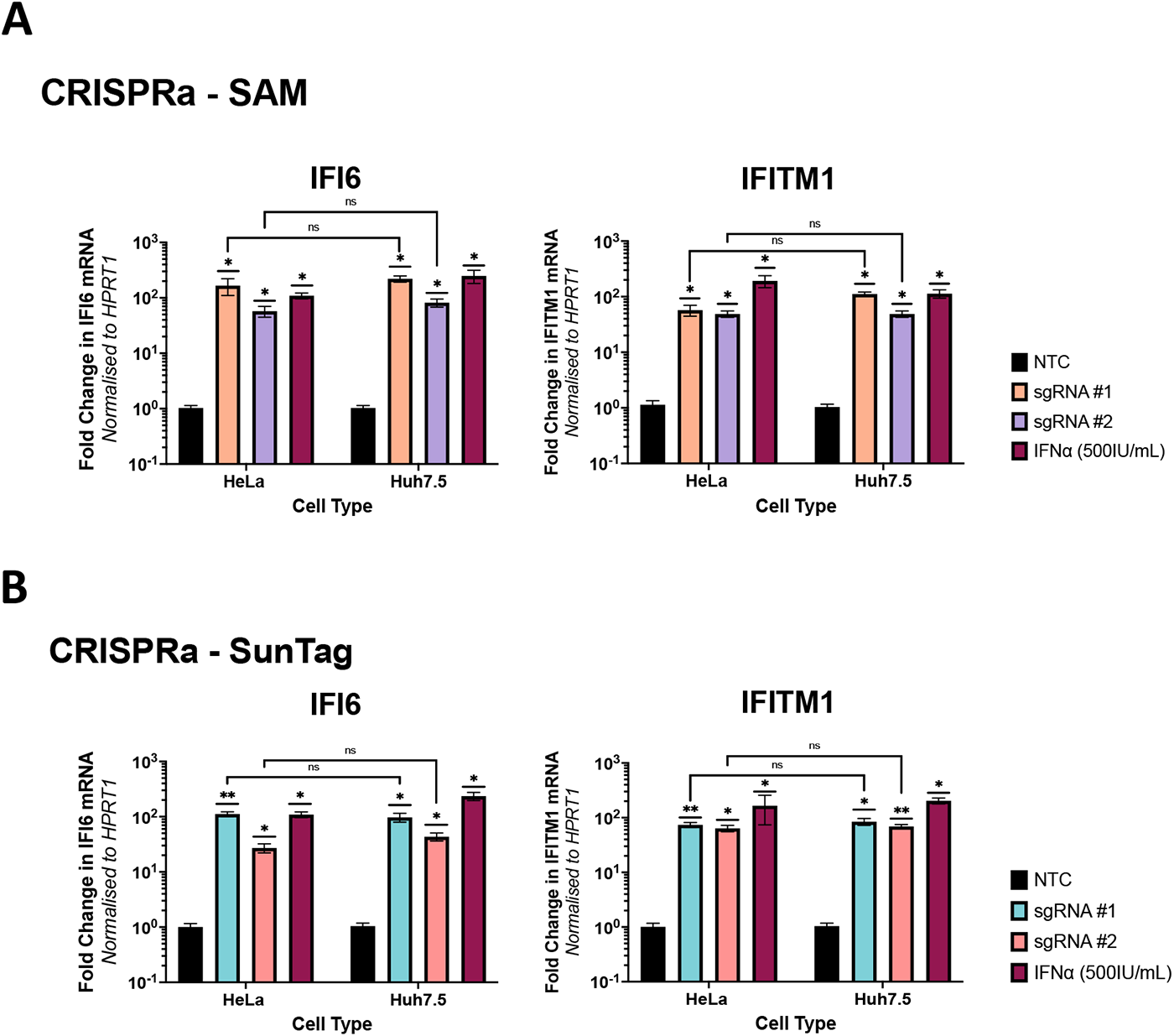
CRISPRa-SAM and SunTag transcriptional activation of ISGs is not significantly different between cell types. HeLa and Huh7.5 cells were transfected with the required plasmids for expression of the CRISPRa-SAM **(A)** or CRISPRa-SunTag **(B)** systems, including sgRNA specific for IFI6 or IFITM1. 72hrs post-transfection RNA was harvested and qRT-PCR analysis was performed to determine if either CRISPRa system induced differing levels of transcriptional activation in HeLa’s versus Huh7.5’s. Statistical analysis was performed by One-Way ANOVA where where * P ≤ 0.05, ** P ≤ 0.01, *** P ≤ 0.001, **** P ≤ 0.0001 (n=3)

### CRISPRa mediated constitutive transcriptional activation of ISGs

Our CRISPRa ISG transient expression approach outlined above may not be useful for long-term ISG anti-viral screening experiments that require rapid scalability. Therefore, we generated Huh7.5 cells constitutively expressing CRISPRa ISGs by transduction of lentiviral packaged pXPR_502 integrated with an ISG specific sgRNA, to allow stable integration of the sgRNA and activator complex into Huh7.5 cells stably expressing dCas9-VP64 (data not shown). We elected to generate Huh7.5 polyclonal CRISPRa lines expressing the antiviral ISGs, IFI6, Viperin, IFITM1, IFIT1, ISG15, ISG20, Mx1, IFNβ and IFNλ2 as they have established antiviral for several Flaviviridae members such as ZIKV, DENV, WNV and the human coronavirus HCoV-229E. qRT-PCR analysis confirmed SAM mediated transcriptional activation of all ISG mRNA, greater than or equal to that of IFNα (500IU/mL) stimulation 24hrs prior to RNA harvest (Fig 4). However, we again observed significant variation in efficacy between sgRNAs, consistent with our results presented previously. Cell lysates from CRISPRa IFITM1 (Fig 5A) and ISG15 (Fig 5B) cells were used in western blotting analysis that revealed significant protein expression in Huh7.5 CRISPRa ISG lines in comparison to no protein expression in CRISPRa NTC Huh7.5 cells. In addition, we confirmed CRISPRa IFNλ2 expression using a transwell assay, validating IFNλ2 secretion that can act in a paracrine manner to stimulating ISG expression in naïve Huh7.5 cells. qRT-PCR of naïve Huh7.5 cells, seeded with a transwell insert containing CRISPRa IFNλ2 cells, showed a significant increase in IFITM1 and IFI6 mRNA compared to naïve Huh7.5 cells seeded with a transwell insert containing CRISPRa NTC cells (Fig 5D). Interestingly, this increase in IFI6 and IFITM1 was comparable to that of exogenous IFNα stimulation, confirming that CRISPRa mediated IFNλ2 expression was active in a similar manner to that of exogenously added IFNα.

**Figure 4:**
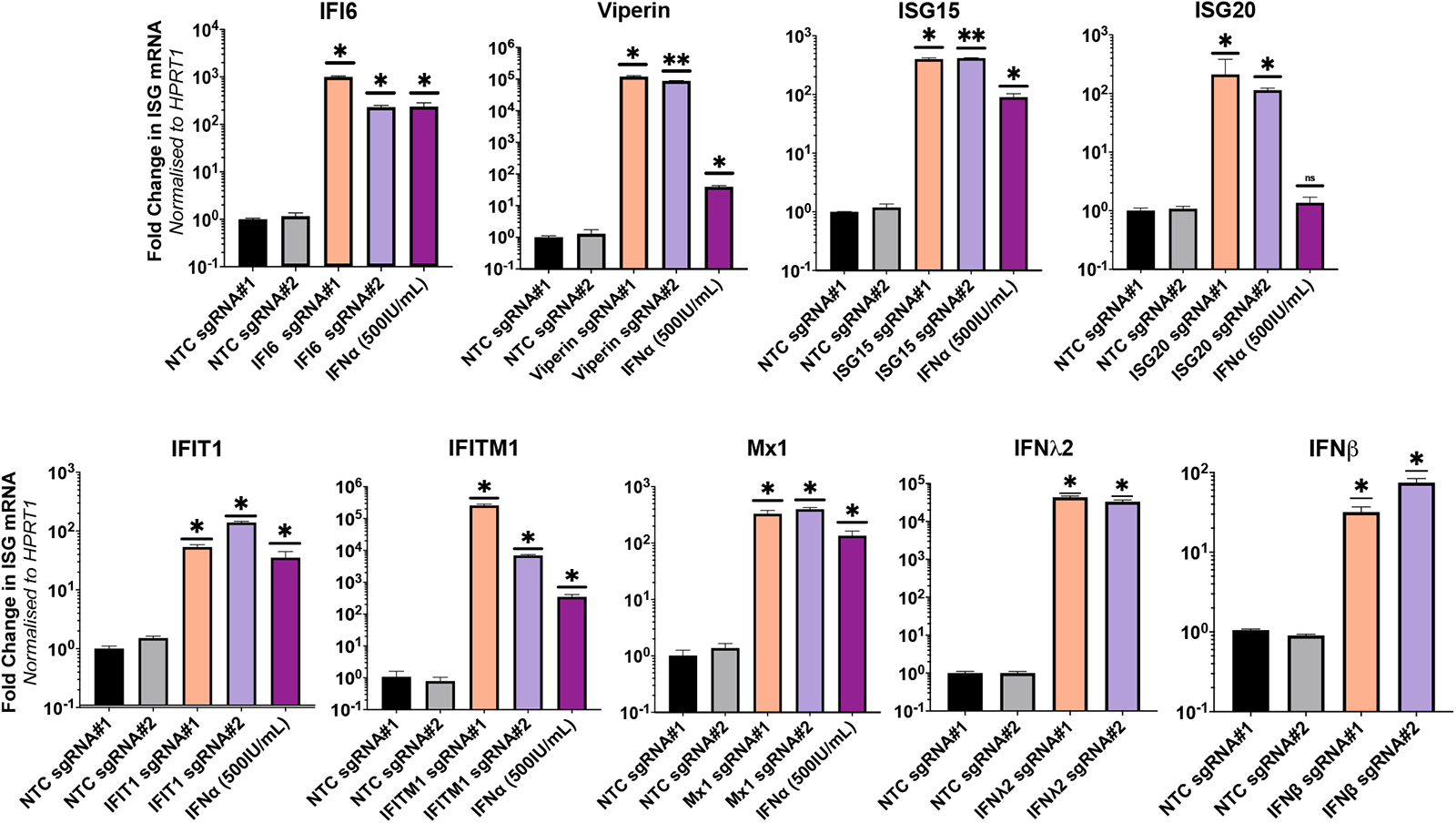
Stable integration of CRISPRa-SAM componentry induces transcriptional activation of target genes. (A) Huh7.5-dCas9 cells were transduced with lentivirus transferring pXPR_502 containing an ISG specific sgRNAs. 72hrs post-transduction, successful integration was selected for with 3µg/mL of puromycin and death of naïve WT Huh7.5 cells, RNA was extracted and qRT-PCR analysis determined the efficacy of CRISPRa mediated ISG transcriptional activation relative to the NTC. Statistical analysis was performed using a One-way ANOVA where * P ≤ 0.05, ** P ≤ 0.01, *** P ≤ 0.001, **** P ≤ 0.0001.

**Figure 5:**
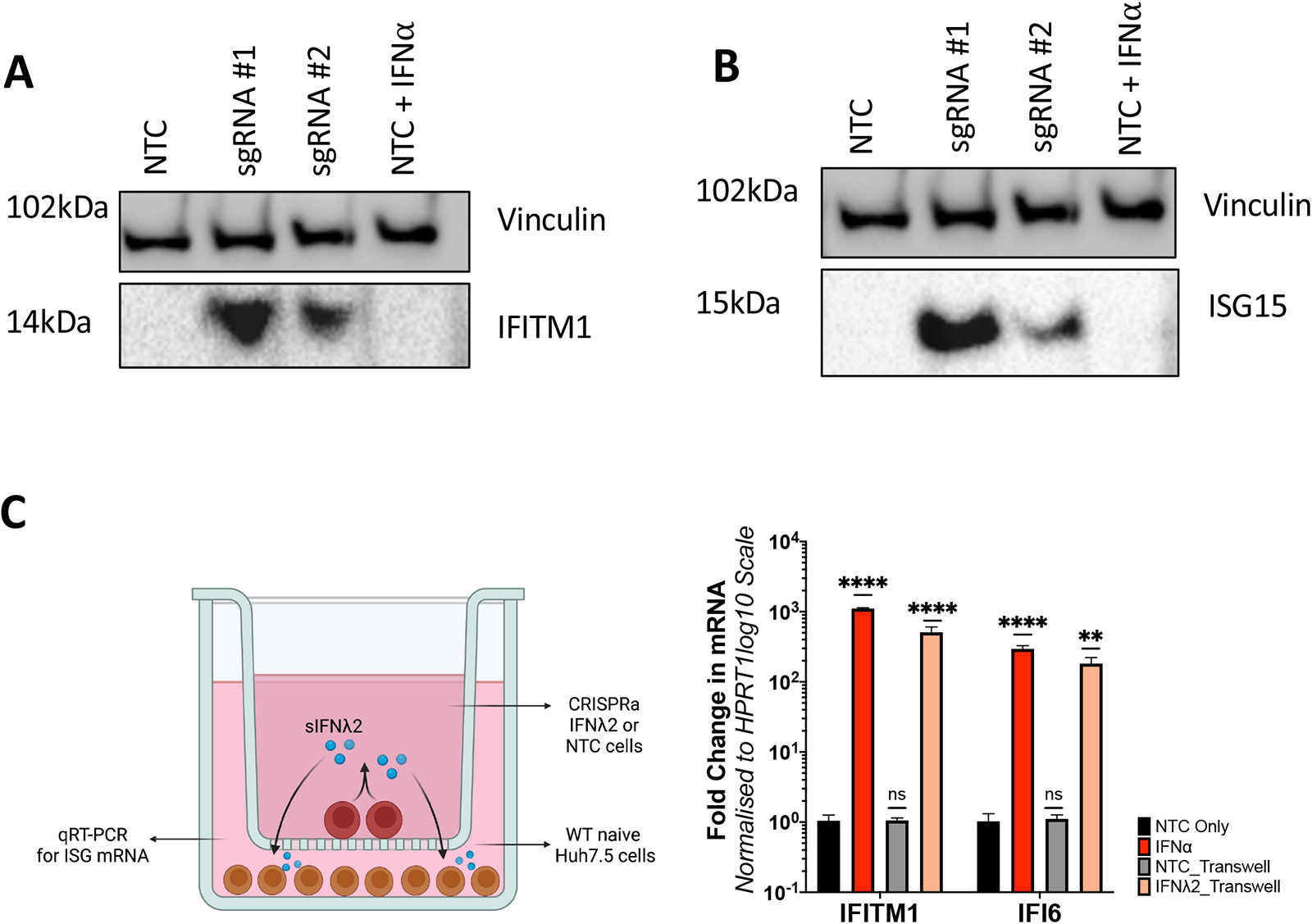
CRISPRa mediated transcriptional activation of select ISGs results in increased protein expression. Lysates of CRISPRa IFITM1 **(A)** and ISG15 **(B)** cells were used in western blotting to confirm transcriptional activation of the target ISG increased relative to CRISPRa NTC cells. **(C)** A transwell assay confirmed IFNƛ2 secretion (sIFNƛ2) from CRISPRa IFNƛ2 where naïve Huh7.5’s were seeded directly into a well, while Huh7.5-IFNƛ2 sgRNA #1 or NTC cells were seeded into a transwell insert. 8hrs post seeding, the insert was layered on top of the well and following 24hrs of culture, total RNA was extracted from the naïve Huh7.5’s and qRT-PCR performed to determine expression of IFI6 or IFITM1. Statistical analysis was performed using a One way ANOVA where * P ≤ 0.05, ** P ≤ 0.01, *** P ≤ 0.001, **** P ≤ 0.0001.

Collectively our qRT-PCR and western blotting indicate that CRISPRa SAM mediated ISG transcriptional activation is an efficient method for ISG expression to levels that we similar or greater than exogenous IFNα stimulation, suggesting that CRISPRa can overcome traditional mechanisms of gene regulation to gene expression. This provides the opportunity to study the anti-viral activity of ISGs independently of the hundreds of ISGs expressed following exogenous IFNα stimulation. It also highlights the utility of CRISPRa ISG expression to be used in the development of arrayed screening platforms to identify and characterise the role of anti-viral ISGs against multiple different viral lifecycles.

### CRISPRa of ISGs can inhibit virus induced cytopathic cell death

To determine if CRISPRa mediated ISG expression can prevent viral induced cytopathic cell death (CPE), CRISPRa ISGs cells were infected with either the flaviviruses, ZIKV (MOI 0.5), WNV_(KUNV)_ (MOI 0.1) or the unrelated coronavirus HCoV-229E (MOI 0.1). Infected cultures were monitored daily for complete CPE of CRISPRa NTC cells, after which cells were formalin fixed and stained with crystal violet (CV) to visualise and quantitate surviving cells in an assay we have termed cytopathic protection assay (CPA). Our CPA identified the ISGs, IFI6, IFNλ2 and IFNβ, that consistent with the literature provided significant protection of cells from WNV_(KUNV)_ and ZIKV induced CPE (Fig 6A and 6B). However, the ISGs IFITM1, that has antiviral potential, provided little cellular protection against WNV_(KUNV)_ and none against ZIKV. These observations were quantitated by measuring retained CV absorbance for each CRISPRa ISG line that supported our visual observations. This highlights the capabilities of the CRISPRa SAM system to characterise the anti-viral potential of ISGs between closely related viruses, that already have a well characterised ISG profile providing an opportunity to identify minute differences that may a yield a better understanding of viral-host interactions. Complete images of CV retention can be found in Fig S1.

**Figure 6:**
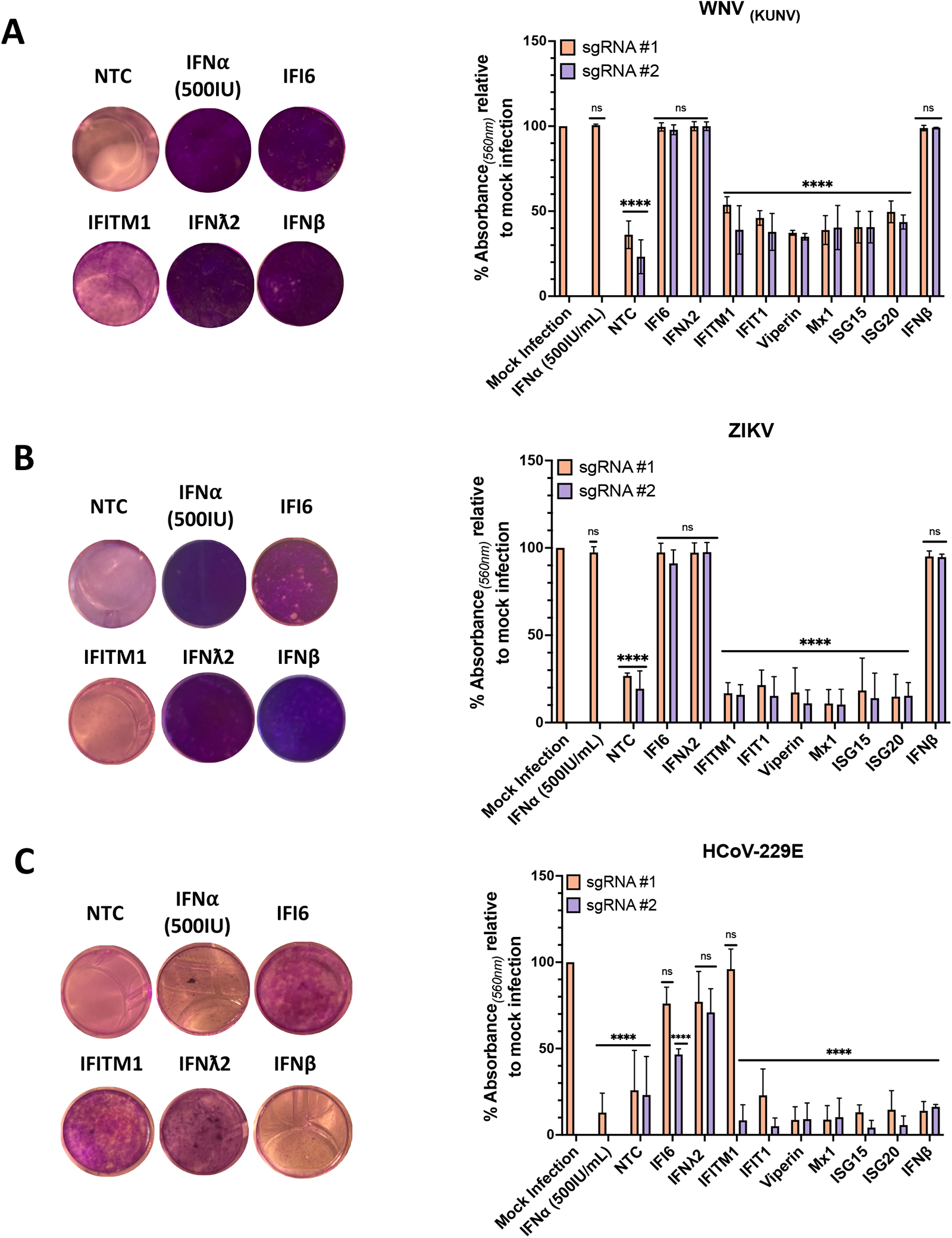
Protection from virally induced cytopathic cell death is ISG dependent. CRISPRa ISG cells, CRISPRa NTC cells or naive Huh7.5 cells stimulated with IFN⍺ (500IU/mL) were infected with **(A)** WNV_(KUNV)_ (MOI 0.5), **(B)** ZIKV (MOI 1) or **(C)** HCoV-229E (MOI 0.1) and observed daily for CPE. Upon complete CPE of CRISPRa NTC cells (ZIKV – 5dpi, WNV_(KUNV)_ and HCoV-229E – 4dpi) wells were fixed with 10% formalin and stained with CV solution. Images were cropped using GIMP photo editing software. CV retention was quantitated by addition of 100% methanol and absorbance (560nm) measured using the Promega GloMax. CV retention for each CRISPRa ISG cell line is expressed as % absorbance relative to mock infected controls. Statistical analysis, compared to the mock infection, was performed using a Two-way ANOVA (n = 3) where ns = non-significant and **** P≤ 0.0001.

To highlight the utility of the CPA across different virus families, we next used the same CRISPRa ISG cells with HCoV-229E. Little is understood of the ISG response for control of HCoV-229E infection. Like ZIKV and WNV_(KUNV)_, we observed potent inhibition of HCoV-229E induced CPE by IFNλ2, in addition to IFI6 and IFITM1 (though only for sgRNA #1) (Fig 6C). Interestingly, no other ISGs prevented CPE, but of particularly note is a lack of anti-viral activity by CRISPRa IFNβ and exogenous stimulation with IFNα (500IU). As both are type I IFNs with near identical signalling pathways, it is plausible that HCoV-229E is capable of silencing type I IFN pathways, preventing downstream activation of an anti-viral state. However, HCoV-229E is still permissive to IFNλ2, a type III IFN (Fig 6C). This observation is in line with recent coronavirus studies highlighting the sensitivity of SARS-CoV-2 to type III IFNs more so than type I, thus warranting further investigation [30]. Again, this highlights the ability of CRISPRa SAM as a tool to characterise the ISG response to infection of less understood viruses.

### CRISPRa SAM identifies ISGs as inhibitors of Hepatitis A Virus replication

The viruses used above all display CPE that provides a useful phenotypic readout for our CRISPRa ISG studies. However, not all viruses induce CPE and thus alternate methods of viral replication must be used. We therefore used Hepatitis A Virus (HAV) (HM175/18f) in which nanoluciferase (NLuc) was inserted between pX and the mature peptide 2B (Fig 7A), termed HAV-NLuc [31–33]. CRISPRa ISG cells were infected with a 0.25 dilution of the generated HAV-NLuc viral stock and cell lysates were collected at the specified time points post-infection. Our understanding of the ISGs that control HAV are not well developed, however our studies suggest that IFI6, IFNλ2, Viperin, IFIT1 (late), Mx1, IFNβ and IFITM1 all have the ability to control HAV replication at least in cultured Huh7.5 cells (Fig 7B). This suggests that ISGs can control HAV infection that may contribute to the acute nature of this infection or alternatively the ability of HAV to efficiently supress the innate antiviral response is a mechanism of overcome ISG expression to allow for viral replication in hepatocytes. Whatever the mechanism further studies are clearly required to determine the repertoire of ISGs that control HAV. Moreover, these results highlight the agile utility of CRISPRa across different viral replication models.

**Figure 7:**
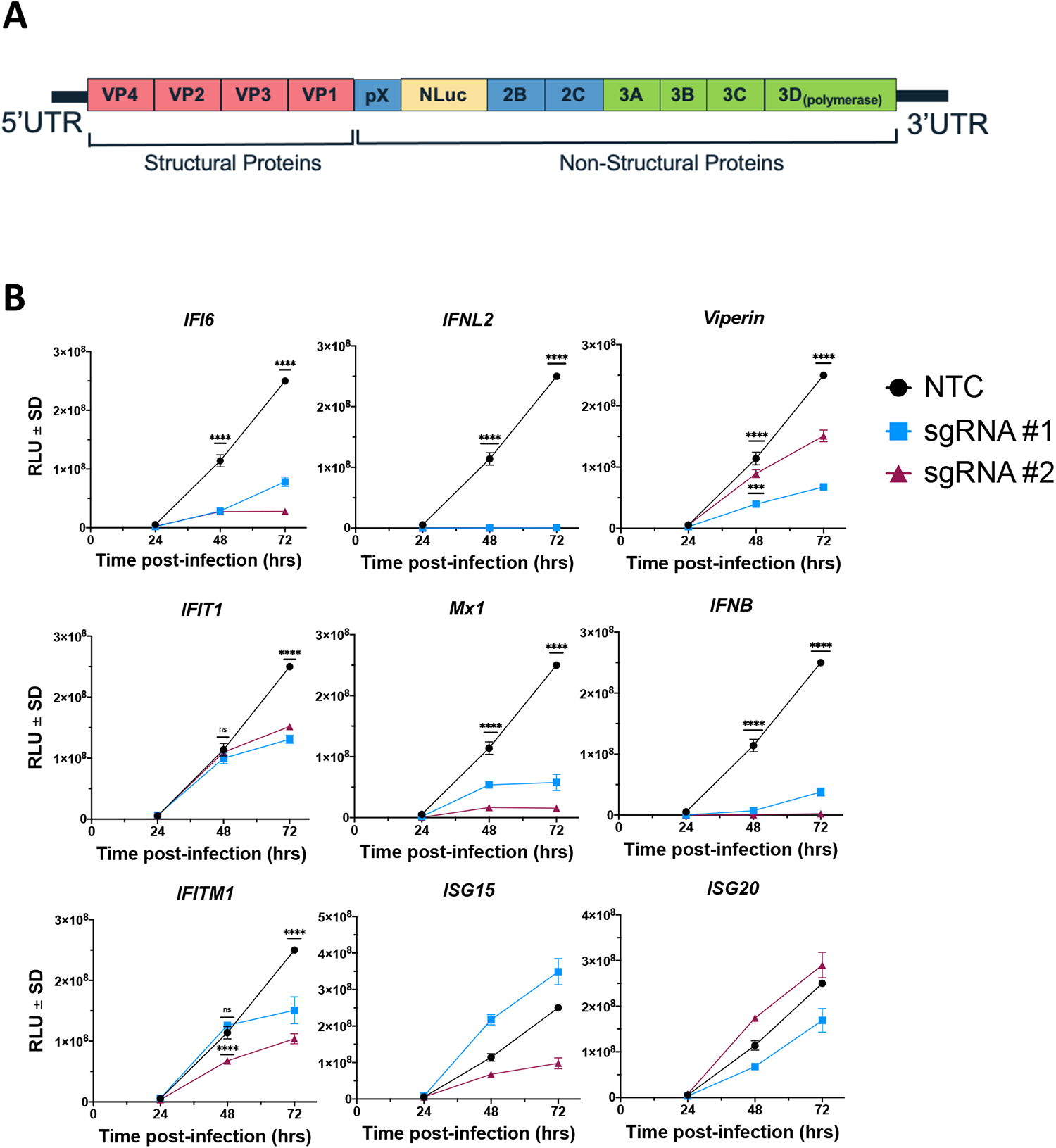
CRISPRa ISG screening identifies inhibitors of HAV-NLuc replication. **(A)** Recombinant HAV-NLuc viral stocks were generated from *in vitro* transcribed RNA of the HM175/18f genome with NLuc inserted between the non-structural proteins pX and 2B as outlined. **(B)** CRISPRa ISG and NTC cells were infected with a ¼ dilution of the generated HAV-NLuc stock. Cell lysates were collected at the specified time points post-infection and a nanoluciferase assay performed to determine HAV replication in response to the presence of the specified ISG. Statistical analysis was performed using a Two-way ANOVA with Tukeys multiple comparisons *** P ≤ 0.001, **** P≤ 0.0001

### Identification of anti-viral ISGs utilising high-content imagining to identify inhibitors of flavivirus replication

Further to our CPA, we developed a high-content imaging (HCI) method to detect and quantify flavivirus replication in an arrayed and unbiased format. HCI provides many benefits as a screening method in that significant amounts of information can be delineated based upon the experimental parameters that can be feasibly upscaled allowing for large scale data collection. Furthermore, HCI can be utilised to overcome the need to use CPE as an endpoint. To develop our HCI screen, CRISPRa ISG cells were seeded into 96-well black plates and infected with either ZIKV, DENV or WNV_(KUNV)_. 48hrs post-infection, fixed wells were subjected to immunofluorescence labelling with an anti-flavivirus envelope antibody (4G2), followed by conjugation with anti-mouse IgG AlexFluoro-488. Cell nuclei were stained with DAPI. Automated imaging was performed using the ArrayScan XTI High Content Analysis Reader (Themo Fisher Scientific). For each well, DAPI staining was used to identify 2000 individual cells as primary objects, followed by quantitation of AlexFluoro-488 signal in the cytoplasmic space surrounding the nucleus. The mean fluorescence intensity (MFI) of total AlexFluoro-488 (488/496nm) was quantified for each well, with three technical replicates per biological replicate used (Fig 8A). Infection is expressed as a % of the MFI of each CRISPRa ISG cell line relative to the MFI of CRISPRa NTC cell for **(B)** DENV, **(C)** ZIKV and **(D)** WNV_(KUNV)_. Furthermore, the efficacy of ISG anti-viral activity was classified as no significant inhibitory activity (80-100% – blue), modest inhibition (51-79% – green) and potent inhibition (0-50% – pink). Additionally, we confirmed the reproducibility of infection across biological replicates by determining that the R^2^ correlation coefficient for each virus was greater than 0.5 (see Table S3 for complete R^2^ values). Based upon the above criteria, HCI identified IFNλ2, IFNβ and IFI6 as potently anti-viral towards all flaviviruses, consistent with our findings from the CPA. Interestingly, this is where shared similarities between all flaviviruses end as our analysis identified virus specific ISGs with modest inhibitory activity such as Viperin (sgRNA#2) towards ZIKV and WNV_(KUNV)._. This highlights that HCI of CRISPRa cells has the capacity to identify both common and virus specific anti-viral ISGs and has validated the known anti-flavivirus activity of several ISGs.

**Figure 8:**
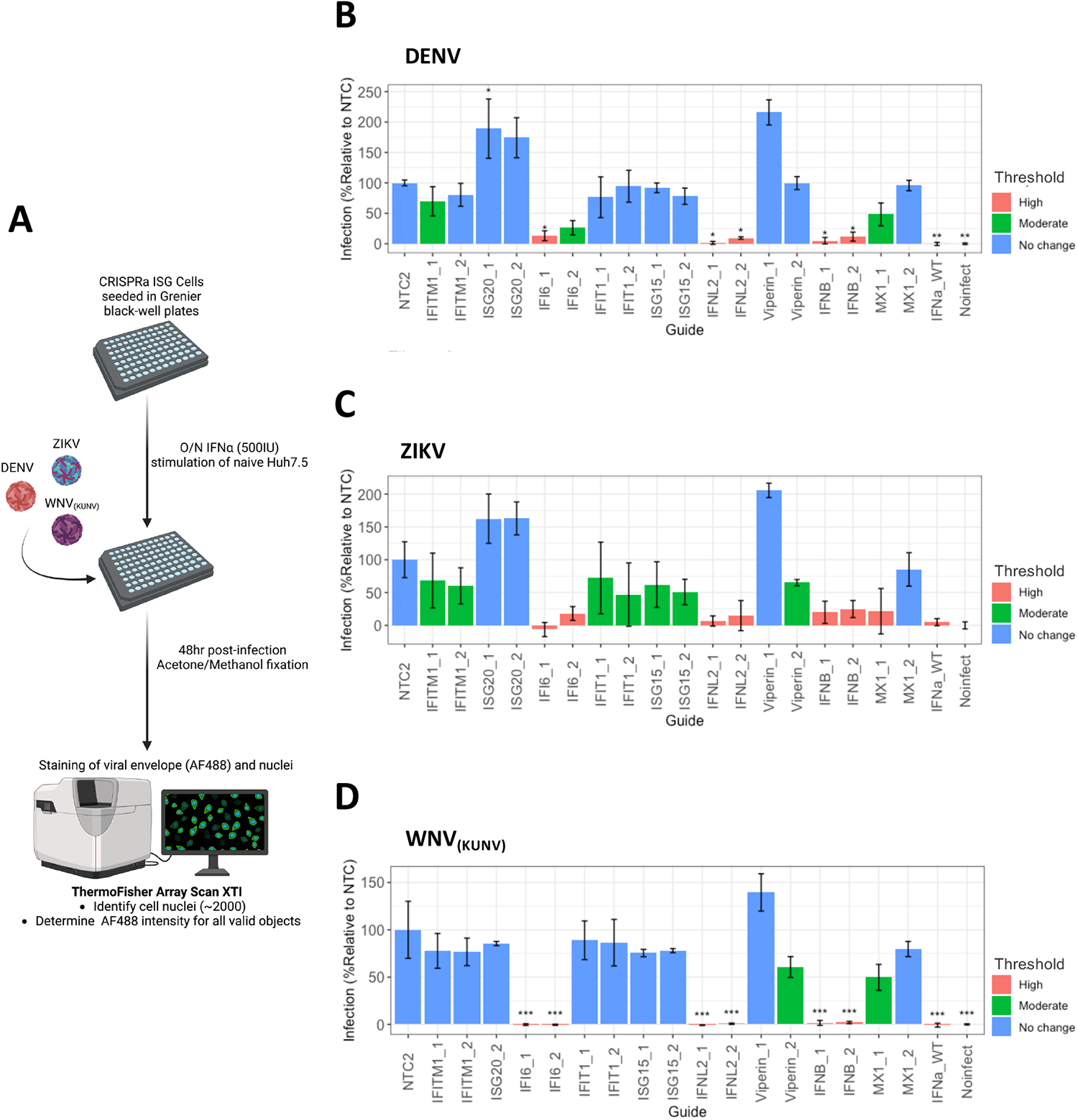
High-content imaging of CRISPRa ISG cells identifies inhibitors of flavivirus replication. **(A)** CRISPRa ISG cells were seeded into black-well plates (Grenier) 24hrs prior to infection with either WNV_(KUNV)_ (MOI 0.5), ZIKV or DENV (MOI 1). 48hrs post-infection, wells were fixed and stained with anti-E (4G2) mouse hybridoma supernatant and AlexaFluro488 secondary (FITC), and nuclei visualised with DAPI. Using the Thermo Fisher XTI Array Scan, wells were imaged to identify ∼2000 primary objects (DAPI) where the FITC ring intensity surrounding each primary object was determined. For each CRISPRa ISG line, the mean ring intensity (MFI) was calculated and expressed as % infection relative to the MFI of CRISPRa NTC for **(B)** DENV, **(C)** ZIKV and **(D)** WNV_(KUNV)_. ISG anti-viral activity was classified based on pre-determined thresholds where % infection between 80-100% is no change (blue), 51-79% is moderate (green) and 0-50% is high (pink) level viral inhibition. Statistical analysis was determined by One-way ANOVA where * P≤0.05, ** P≤0.01 and *** P≤0.001 (n=3).

Collectively, our results highlight the utility of CRISPRa mediated ISG expression in routine and high-throughput formats to delineate the role of ISGs on inhibition of viral replication.

## Discussion

The production of ISGs by the host innate response to viral infection is critical to controlling viral replication and dissemination [34]. Understanding the complex interplay between virus and host is crucial to enhancing our knowledge of viral pathogenesis, disease severity and the possibility of anti-viral therapeutic development. Concerted efforts have been made to characterise the anti-viral mechanism of the ∼ 300 known ISGs, which have generally been investigated as a whole in response to a stimulus (e.g. viral infection or exogenous IFN stimulation) [35,36]. Attempts to characterise ISGs individually, *in vivo* and *in vitro*, have been achieved through the use of gene editing technologies like RNA interference (RNAi – siRNA, shRNA) or CRISPR_KO_, where the impact of the ISG is determined by alterations in the rate of viral replication [7,37–41]. Alternatively, ectopic overexpression has proven successful, allowing individual ISGs to be expressed independently from the canonical IFN signalling pathway [42,43]. However, the traditional silencing and overexpression mechanisms come with limitations. cDNA overexpression screens typically produce excessive levels of the target ISG, which maybe not physiologically relevant for the cell type in which it is expressed, nor does it consider post-translational modifications such as splice variants. Furthermore, siRNA and cDNA methods have poor scalability, in that both are costly and time-expensive [44]. Additionally, the result of identified anti-viral host factors is often divergent between RNAi and CRISPR_KO_ screens, where the lack of consistency makes it difficult to define what is truly anti-viral and what is not [39]. CRISPRa has emerged as an alternative method due to its ease to scale appropriately and comparatively low cost [44]. CRISPRa strongest benefit lies it is ability to induce endogenous gene activation from the target promoter, where effective sgRNA design allows transcriptional activation with the aim to achieve the “static upper limit”, the point of physiologically relevant maximum expression [16,17,45]. Endogenous activation also allows for the expression of all transcript variants. The commercial availability of genome-wide CRISPRa screening sgRNA libraries has allowed for the rapid identification of anti-viral host factors for several viruses like IAV, ZIKV and SARS-CoV-2 [21–23,41]. However, many of these host factors were not ISGs, which given their potency to inhibit viral replication, the question was raised, is CRISPRa a suitable method to study ISG function in various screening formats. It is plausible that CRISPRa induction of ISGs does successfully inhibit viral replication, but the nature of genome-wide screening, depending on the endpoint (e.g. CPE), generally identifies host factors with the strongest phenotypic inhibition [46]. This has been observed in several publications where the ISG IFI6 was identified as anti-ZIKV, while OAS1 was anti-SARS-CoV-2 [21,22]. Therefore, we set out to develop and characterise several arrayed screening platforms to determine if CRISPRa has the potential to identify both strong and modestly anti-viral ISGs.

With this in mind, we firstly sought to determine the capability of two CRISPRa systems, SAM and SunTag, to induce ISG expression. Transfection of the respective system vectors resulted in SAM consistently outperforming SunTag at inducing IFI6 and IFITM1 mRNA expression both overtime (Fig. 2) and in different cell types, HeLa and Huh7.5’s (Fig. 3). These differences likely arise from several mechanistic differences between the systems, such as the need for CRISPRa-SunTag to express VP64 ten times that of CRISPRa-SAM to ensure maximal recruitment of host transcription factors [16]. A key observation from this work is the significant differences in efficacy between the sgRNAs for each target ISG. For example, we noted a ∼0.5 log_10_ fold change between IFI6 sgRNA #1 and sgRNA #2 (Fig. 3A and 3B) even though they are positioned within 100bp of each other. This variation continues upon generating constitutively expressing CRISPRa-SAM ISG lines, though mRNA levels are significantly increased due to stable integration of the CRISPRa-SAM componentry (Fig. 4). This highlights a critical, continuing limitation of all CRISPRa systems, sgRNA design. Computational programs used in sgRNA design, such as FANTOM, are a continually developing field that will lead to more effective sgRNA design and enhanced transcriptional activation [16,17]. In the meantime, it is imperative to develop scalable screening methods that can easily be adapted according to experimental requirements. This includes the use of multiple target gene sgRNA and the use of constitutively activated lines, allowing for rapid screening to avoid the introduction of experimental error that may further alter the observed phenotype.

Using a CRISPRa ISG cell panel, we developed several routine and HCI arrayed screening platforms to assess the utility of our approach and confirm ISG anti-viral activity against the flaviviruses WNV_(KUNV)_, DENV and ZIKV and the unrelated HCoV-229E. Firstly, we performed a series of viability assays to determine if our CRISPRa ISG panel was capable of preventing viral induced CPE. Unsurprisingly, IFI6 was a potent inhibitor for the flaviviruses, but also provided some protection against HCoV-229E induced CPE (Fig. 6). This potent anti-flavivirus activity of IFI6 in our CPA was validated by our HCI approach (Fig 8C). It is well established that IFI6 inhibits flavivirus replication by preventing formation of replication complexes, however, IFI6 has not been shown to inhibit α-coronaviruses HCoV-OC43 or the β-coronaviruses HCoV-NL63 [7] . It is plausible that HCoV-229E is sensitive to IFI6, as coronaviruses also induce alterations of ER membranes for formation of replication complexes, however, it is more likely that as IFI6 has anti-apoptotic properties, that the surviving cells are expressing IFI6 to an acceptable level to prevent CPE [47,48]. IFITM1, particularly sgRNA #1, was also a standout from our viability panel, displaying enhanced anti-viral activity towards HCoV-229E, and modest inhibition of WNV_(KUNV)_, but not ZIKV, an interesting observation given that IFITM1 has been reported to be anti-ZIKV using both shRNA and ectopic overexpression methods [49]. Meanwhile, the anti-coronavirus activity of IFITM1 has been well established as IFITM1 is potent inhibitor of non-TMPRSS2 mediated coronavirus entry [50,51]. Overall, this suggests that a combinatorial approach to ISG screening is required as each methodology has advantages and disadvantages, however CRISPRa may provide a more physiologically relevant insight given transcription is driven from the endogenous promoter.

Unexpectedly in our CPA, Viperin did not protect against flavivirus induced CPE, though its anti-viral effects are well established, be it through direct inhibition of the viral non-structural proteins or augmentation of PRR signalling in response to virus detection [52–54]. However, in analysis of our HCI screen, a more direct measurement of genome replication and translation, Viperin sgRNA #2 was a potent inhibitor of ZIKV and WNV_(KUNV)_ (Fig 8). The contrast between the two screening methods highlights that upon further optimisation, CRISPRa is capable of further delineating an ISGs anti-viral mechanism by determining the efficacy of inhibition throughout various stages of the viral lifecycle. In the case of Viperin, the majority of flavivirus genome replication is likely being inhibited, however, over the period of several days, the compounding effect of some virus survival overwhelms the anti-viral capacity of Viperin, promoting virally induced CPE. Furthermore, Mx1 (sgRNA #1) was identified as an inhibitor (though not significant) of flavivirus infection (Fig 8). Mx1 has not been considered a crucial inhibitor of flavivirus replication as previous studies with Japanese Encephalitis Virus, Hepatitis C Virus and WNV_(KUNV)_ have not been affected by ectopic overexpression of Mx1, as it is hypothesised that progeny genomes are protected from Mx1 activity by establishment of the replication complex [11,55–57]. Firstly, we must consider that CRISPRa Mx1 expression may be driving Mx1 expression that is significantly greater than IFN stimulation, leading to the inadvertent localisation of Mx1 to the ER, preventing virion formation. Alternatively, Mx1 has ∼48 transcript variants, therefore making the antiviral effect of Mx1 difficult to assess when using cDNA ectopic overexpression [58]. The inhibitory activity of Mx1 in our HCI may be due to CRISPRa induction of transcription from the endogenous promoter, allowing for the expression of all Mx1 transcript variants which is extremely difficult using ectopic overexpression methods. Furthermore, given the success of our arrayed screens to identify anti-flavivirus and coronavirus ISGs, we expanded our studies to identify antiviral ISGs of HAV infection for which data is lacking. Interestingly in contrast to the flaviviruses our screen identified that the majority CRISPRa ISGs investigated are anti-HAV with the exception of ISG15 and ISG20 (Fig. 7). Clearly future studies are required to determine the complete repertoire of ISGs that can control the HAV life cycle and how this impacts infection outcome.

The use of ISG CRISPRa expression now provides a framework to identify anti-viral ISGs for HAV and other viruses of interest. Therefore, our arrayed screens to identify anti-viral ISGs against well characterised viruses has identified both known and novel insights into the mechanism of anti-viral replication, while proving that CRISPRa is an appropriate tool to study the viral-host relationship.

In summary, our CRISPRa ISG anti-viral screening platforms confirmed the anti-viral activity of a number of well-characterised ISGs and identified novel inhibitors of viral replication. While we have demonstrated the utility of CRISPRa technology to identify the anti-viral activity a select group of ISGs independently of activation of the innate response, there are a number of limitations. Firstly, we must consider the synergistic effect that ISGs may possess when expressed in combination. A prime example of this is the relationship between the ISGs Viperin and CMPK2 (a monophosphate kinase) in which previous work has indicated that Viperin is capable of converting dCTP to ddhCTP, a ribonucleoside analogue acting as a chain terminator of DENV, WNV, ZIKV and HCV genome replication [59]. When Viperin and CMPK2 are co-expressed, the ratio of dCTP to ddhCTP conversion significantly increases overtime compared to Viperin expression alone, suggesting that CMPK2 provides a constant source of dCTP for Viperin [59]. Future experiments that aim to characterise the synergistic effects of ISGs should utilise bioinformatics pipelines to identify ISGs that share promoters or other ISG regulatory elements to delineate possible CRISPRa sgRNA multiplexing formats. Furthermore, a number of ISGs are toxic when constitutively expressed (a reason why IFN responses are tightly regulated) and thus future experimental CRISPRa platforms should incorporate inducible CRISPRa or regulated ISG expression. Nevertheless, CRISPRa technology has significant potential to characterise the repertoire of ISGs that control viral replication across a broad range of medically relevant viruses.

## Acknowledgements

This work was supported by the NHMRC of Australia (APP2004090) and The University of Adelaide Biochemistry Trust Fund. Emily N. Kirby, Xavier. B Montin, Timothy P. Allen and Brooke N. Trowbridge would like to acknowledge the support of The University of Adelaide Research Training Program Stipend (RTPS). The authors would like to acknowledge the facilities provided by Adelaide Microscopy, and Dr. Agatha Labrindis for providing technical assistance.

## Author contributions

Emily N. Kirby, Xavier B. Montin, Timothy P. Allen, Jaslan Densumite, Brooke N. Trowbridge conducted experiments and analysed the results. Emily N. Kirby, Timothy P. Allen and Michael R. Beard provided experimental design input. Michael R. Beard acquired funding. Emily N. Kirby and Michael R. Beard conceived the study and wrote the manuscript.

## Materials and Methods

### Cell and culture conditions

All mammalian cell lines were maintained at 37°c in a 5% CO_2_ air atmosphere. Huh7.5 human hepatoma cells, HEK293T human embryonic kidney cells and Vero African Green monkey cells were maintained in DMEM (Gibco) containing 10% (v/v) FCS and 1% (v/v) penicillin and streptomycin. MRC-5 human fetal lung fibroblasts were maintained in RPMI (Gibco) containing 10% (v/v) FCS and 1% (v/v) penicillin and streptomycin. C6/36 *Aedes albopictus* cells were maintained in Basal Medium Eagle (Gibco) supplemented with L-glutamine, MEM non-essential amino acids, sodium pyruvate, 10% FC and P/S and culture at 28°c in a 5% CO_2_ air atmosphere.

### Antibodies and Chemicals

Mouse anti-envelope glycoprotein 4G2 (D1-4G2-4-15) was prepared from hybridoma cells purchased from ATCC. Mouse anti-IFITM1 was purchased from ProteinTech (60074-1-Ig) and Rabbit anti-ISG15 was purchased from Abcam (ab133346). Secondary AlexaFluor 488 (A-11029), goat anti-mouse IgG HRP (A24512) and DAPI (D1306) were purchased from Thermo Fisher Scientific. Goat anti-Rabbit IgG HRP (GERPN4301) was purchased from Sigma Aldrich.

### Viruses and Plasmids

ZIKV PRVABC59 (Puerto Rico, 2015) was originally obtained from ATCC. WNV_(KUNV)_ (NSW2011) was generously donated by Karla J. Helbig (La Trobe University, Melbourne, Australia), HCoV-229E was kindly provided by Natalie A. Prow (University of South Australia, Adelaide, Australia) and HAV-NLuc (HM175/18f) was kindly supplied by Stanley M. Lemon (University of North Carolina, Chapel Hill, USA)[33]. DENV2 (strain 16681) was generated from plasmid pFK-DVs containing the full-length genome, generously donated by Ralf Bartenschlager (University of Heidelberg, Heidelberg, Germany), as described in [60]. Infectious viruses were amplified in either C6/36 cells (ZIKV, DENV and WNV_(KUNV)_), MRC-5 cells (HCoV-229E), clarified by centrifugation at 500 x g for 10 min at 4°c, aliquoted and stored at -80°c. Virus titre was determined by plaque assay or focus-forming assay (DENV) as described in [60].

### Generation of Huh7.5 Stable ISG Lines

To generate stably ISG expressing CRISPRa lines, 20nt guide sequences targeting the specified ISGs were selected from the Calabrese CRISPRa sgRNA library and synthesised (IDT), annealed and ligated into *Bsm*BI digested pXPR_502. Following the generation of lentiviruses carrying pXPR_502, monoclonal Huh7.5 cells expressing dCas9 were transduced and puromycin selected for 3 days post transduction until complete death of WT Huh7.5s was observed. Surviving CRISPRa ISG cells were validated by qRT-PCR for transcriptional activation, prior to being used in infection experiments as specified.

### High content screening for viral detection

Stable CRISPRa ISG Huh7.5 cells were seeded at 1 x 10^4^ cells/well into 96-blackwell plates and cultured for 24hrs prior to infection with the specified viruses. Plates were cultured for an additional 48hrs prior to removal of the media and washed 2x with PBS. Cells were fixed with ice-cold acetone:methanol (1:1) at 4°c for 10 min. Following removal of the fixative and washing with PBS, fixed monolayers were blocked with 5% bovine serum albumin (BSA) in PBS at room temperature for 30 min. BSA was then removed and anti-E hybridoma cell supernatant diluted 1:5 in PBS was added and incubated at 4°c for 1hr. Following PBS wash, monolayers were incubated with Alexa Fluor-488 conjugated anti-mouse IgG (Life Technologies) for 1hr at room temperature, followed by PBS washing and incubation with DAPI (4’,6-diamidino-2-phenylindole dihydrochloride [Sigma-Aldrich]) for 10 min at room temperature. Automated imaging was performed using the ArrayScan XTI High Content Analysis Reader (Themo Fisher Scientific). Briefly, cell populations were imaged by the 10x magnification with widefield lenses. Individual cells were identified by DAPI staining (386/23nm) and nuclear segmentation with isodata thresholding. Ring projections of 14 pixels was used to define the cytoplasmic space around each nucleus for EGFP quantification. Total GFP (485/20nm) MFI was quantified for each cytoplasmic ring projection, excluding overlapping areas. 2000 individual cells were quantified per well, with three technical replicates per biological replicate used. Data was background subtracted with the no infection negative control and relative infection was compared to NTC control group. Quantification was done with HCS Studio^TM^ Cell Analysis Software (Thermofisher) and data was analysed with R (version 4.2.2).

### Quantitative Real Time PCR

Cells were seeded in 24 well plates 24hrs prior to infection or transfection. Total RNA was extracted from cells using NucleoZOL (Macherey Nagel) as per the manufacturers instructions. cDNA synthesis and qRT-PCR were performed simultaneously using the Luna Universal One-Step RT-qPCR kit (NEB) with the Quantstudio 7 Flex (Life Technologies) to quantitate relative levels of transcriptional activation or viral RNA compared to the house keeping gene HPRT1. All primer sequences used are outlined in Table S1.

### Western Blotting

Immunoblotting of ISGs was performed using Tris-Tricine SDS systems. Briefly, cell monolayers were washed with PBS and lysed in ice-cold NP-40 lysis buffer (1% NP-40, 150mM NaCl, 50mM Tris pH 8.0) with protease inhibitor cocktail (Sigma-Aldrich). Samples were clarified by centrifugation (10,000 x g, 5 min at 4°c). Each sample was then combined with 2x Tricine loading buffer (0.2M Tris pH 6.8, 40% glycerol, 2% SDS, 0.04% Coomassie blue, 2% β-mercaptoethanol) and boiled at 100°c for 5 min. Samples were separated by SDS-PAGE using Tris-Tricine gels and Tricine running buffer (recipe’s obtained from Proteintech and Phiel Lab respectively [61]) and transferred to Hybond ECL nitrocellulose membrane (GE Healthcare). Following blocking for 1hr at room temperature in TBS-Tween 20 (0.1%) containing 5% skim milk, membranes were incubated overnight with primary antibody in TBST-T containing 1% skim milk. Membranes were then washed in TBS-T (3 x 10 min) and incubated for 1 hr at room temperature with HRP-conjugated anti-mouse or anti-rabbit IgG secondary antibodies diluted in TBS-T containing 1% skim milk. Following additional washing with TBS-T, membranes were developed using SuperSignal West Femto Maximum Sensitivity Substrate (Thermo Fisher Scientific) and image using a ChemiDoc MP imaging system (Bio-Rad).

### HAV-Nano Luciferase assay

Luciferase assays to quantify HAV nano-luciferase (NLuc) levels were performed by seeding of Huh7.5 cells into 48 well plates prior to infection with HAV-NLuc at the indicated dilution. Samples were harvested at specified timepoints with 1x passive lysis buffer before measurement of NLuc activity using the Nano-glo luciferase assay system (Promega) with the GloMax-96 luminometer (Promega).

