## Supplementary Figures for "The utility of CRISPR activation as a platform to identify interferon stimulated genes with anti-viral function"

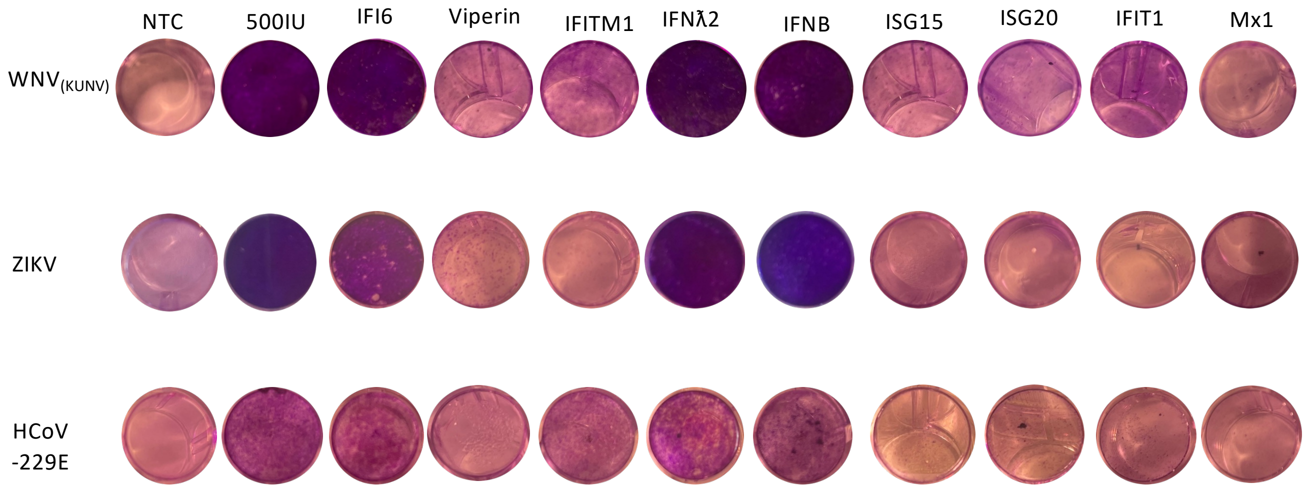


**Figure S1: Full panel of images representing CRISPRa ISG cells following the completion of the CPA**

Images were taken and edited as previously described in Figure 6.

**Table S1: qRT-PCR Primers**

| Gene Name | Primer Forward (5' – 3') | Primer Reverse (5' – 3') |
|---|---|---|
| IFI6 | CTGAAGATTGCTTCTCTTCTC | CACTTTTTCTTACCTGCCTC |
| Viperin | GTGAGCAATGGAAGCCTGATC | GCTGTCACAGGAGATAGCGAGAA |
| IFITM1 | CGCCAAGTGCCTGAACATCT | CCCGTTTTTCCTGTATTATCTGTA |
| IFN$\lambda$2 | ACATAGCCAGTTCAAGTC | GACTCTTCTAAGGCATCTTTG |
| IFN$\beta$ | GCAGTCTGCACCTGAAAAGATAT | TGTACTCCTTGGCCTTGAGGTA |
| IFIT1 | AACTTAATGCAGGAAGAACATGACAA | CTGCCAGTCTGCCCATGTG |
| Mx1 | CAGCACCTGATGGCCTATCAC | CATGAAGAACTGGATGATCAAAGG |
| ISG15 | TGGCGGGCAACGAATT | GGGTGATCTGCGCCTTCA |
| ISG20 | TCGTTGCAGCCTCGTGAAC | TCCCTCAGGCCGGATGA |

### Table S2: CRISPRa sgRNA sequences

| Gene Name | Top Strand (5' – 3') | Bottom Strand (5' – 3') |
|---|---|---|
| IFI6 #1 | AGCACACAAATGTTCCCGCT | AGCGGGAACATTTGTGTGCT |
| IFI6 #2 | AAGTGCATTTTTCTGCCAGC | GCTGGCAGAAAAATGCACTT |
| IFNλ2 #1 | CACAGCCTCAGGTAAGACAC | GTGTCTTACCTGAGGCTGTG |
| IFNλ2 #2 | CAGAGCAGGTGGAATCCTCC | GGAGGATTCCACCTGCTCTG |
| Viperin #1 | GTTATAATACAAGTGGACTG | CAGTCCACTTGTATTATAAC |
| Viperin #2 | AGACTGAAGACTAGAGATCG | CGATCTCTAGTCTTCAGTCT |
| IFITM1 #1 | GGAGGAAAGGCTGAAGGCTA | TAGCCTTCAGCCTTTCCTCC |
| IFITM1 #2 | GGGCCCTGGGGATTTTACCC | GGGTAAAATCCCCAGGGCCC |
| IFNβ #1 | ATGGTCCTCTCTCTATTCAG | CTGAATAGAGAGAGGACCAT |
| IFNβ #2 | AAATTCCTCTGAATAGAGAG | CTCTCTATTCAGAGGAATTT |
| IFIT1#1 | GTGAATCTCGTTCCAAATGC | GCATTTGGAACGAGATTCAC |
| IFIT1#2 | TGCTGCAGAAACCCAATGAC | GTCATTGGGTTTCTGCAGCA |
| Mx1 #1 | AGGCAAGTGCTGCAGGTGCG | CGCACCTGCAGCACTTGCCT |
| Mx1 #2 | CAGAAACGAAACCTAGCTCC | GGAGCTAGGTTTCGTTTCTG |
| ISG15 #1 | GCTCACTCTGGGGCATGCCT | AGGCATGCCCCAGAGTGAGC |
| ISG15 #2 | TCTGGGGCATGCCTCGGGAA | TTCCCGAGGCATGCCCCAGA |
| ISG20 #1 | GCTACCAGTCTGATACTCAG | CTGAGTATCAGACTGGTAGC |
| ISG20 #2 | CATCCCCAGGACTGGAGCTC | GAGCTCCAGTCCTGGGGATG |
| NTC #1 | GTGTCGTGATGCGTAGACGG | CCGTCTACGCATCACGACACC |
| NTC #2 | CATTAGCAGCCCAGCGCCCA | TGGGCGCTGGGCTGCTAATG |

### Table S3: $R^2$ coefficients to determine reproducibility of infection for HCI replicates

| DENV | | | |
|---|---|---|---|
| | **R1** | **R2** | **R3** |
| **R1** | 1.0000000 | 0.8513768 | 0.7076396 |
| **R2** | 0.8513768 | 1.0000000 | 0.8195575 |
| **R3** | 0.7076396 | 0.8195575 | 1.0000000 |

| ZIKV | | | |
|---|---|---|---|
| | **R1** | **R2** | **R3** |
| **R1** | 1.0000000 | 0.5562753 | 0.4053427 |
| **R2** | 0.5562753 | 1.0000000 | 0.8695403 |
| **R3** | 0.4053427 | 0.8695403 | 1.0000000 |

| $WNV_{(KUNV)}$ | | | |
|---|---|---|---|
| | **R1** | **R2** | **R3** |
| **R1** | 1.0000000 | 0.6864971 | 0.8461597 |
| **R2** | 0.6864971 | 1.0000000 | 0.9093724 |
| **R3** | 0.8461597 | 0.9093724 | 1.0000000 |
